# CellAwareGNN: Single-Cell Enhanced Knowledge Graph Foundation Model for Drug Indication Prediction

**DOI:** 10.64898/2026.02.20.707076

**Authors:** Xinmeng Zhang, Eugene Jeong, Chao Yan, Yubo Feng, Linshuoshuo Lyu, Xingyi Guo, You Chen

**Author notes:** Indicates equal contribution (Xinmeng Zhang, Eugene Jeong).

## Abstract

Graph foundation models have emerged as powerful tools for drug repurposing by enabling the prediction of novel drug–disease indications from large biomedical knowledge graphs. A representative example is TxGNN, which was previously developed and trained on PrimeKG, a comprehensive biomedical knowledge graph covering over 17,000 diseases. While TxGNN demonstrates strong performance, existing biomedical knowledge graphs largely lack fine-grained, cell-type-specific expression context. This limits their ability to capture disease mechanisms driven by dysregulated cellular programs, such as immune cell-specific pathways in autoimmune diseases. Moreover, prior evaluations typically test only randomly selected subsets of diseases, leaving many diseases unexamined and limiting conclusions about model performance across the full disease spectrum.

To address these limitations, we first update PrimeKG to **PrimeKG-U** by incorporating expanded and curated biomedical knowledge and then develop **TxGNN-U** as a stronger graph-based baseline. Building on this foundation, we introduce **CellAwareGNN**, a graph foundation model that integrates single-cell genomics into PrimeKG-U. We construct a single-cell-enhanced knowledge graph, **scPrimeKG**, by incorporating cell-type-specific gene expression signatures from the OneK1K dataset, expanding PrimeKG from approximately 8.1 million edges and 129k nodes to over 14 million edges and 147k nodes. CellAwareGNN is pre-trained on all relation types in scPrimeKG and evaluated on drug indication prediction with explicit coverage of all diseases in the knowledge graph.

CellAwareGNN consistently outperforms TxGNN and TxGNN-U. For drug indication prediction, CellAwareGNN achieves an AUPRC of 0.826, representing a 1.2% improvement over TxGNN-U (0.816) and a 3.4% improvement over TxGNN (0.799). Notably, for autoimmune diseases, CellAwareGNN attains an AUPRC of 0.864, improving by 2.0% over TxGNN-U (0.847) and 6.0% over TxGNN (0.815). Importantly, CellAwareGNN prioritizes promising repurposing candidates, including Ocrelizumab for Pemphigus via CD20-expressing B cells, Methotrexate for Pemphigus through DHFR and ATIC activity in T and B cells, and Rosiglitazone for Rheumatoid Arthritis through PPAR-γ activation. These results demonstrate the value of incorporating cell-type-specific expression context to improve both predictive performance and biological interpretability in graph-based drug repurposing.

**CCS Concepts:**

- Applied computing → Health informatics; Bioinformatics;
- Computing methodologies → Knowledge representation and reasoning; *Neural networks*.

**ACM Reference Format:** Xinmeng Zhang, Eugene Jeong, Chao Yan, Yubo Feng, Linshuoshuo Lyu, Xingyi Guo, and You Chen. 2026. CellAwareGNN: Single-Cell Enhanced Knowledge Graph Foundation Model for Drug Indication Prediction. In *Proceedings of the 32nd ACM SIGKDD Conference on Knowledge Discovery and Data Mining V*.*2 (KDD 2026), August 9–13, 2026, Jeju Island, Republic of Korea*. ACM, New York, NY, USA, 12 pages. https://doi.org/10.1145/3770855.3819012

## 1 Introduction

Drug repurposing identifies new therapeutic uses for existing drugs and accelerates treatment discovery [28, 40, 59]. Computational approaches built on biomedical knowledge graphs (KGs) have shown success in drug repurposing by identifying novel drug–disease (indication) pairs [18, 44]. By integrating heterogeneous biomedical data, including genetic factors, molecular interactions, phenotypes, and clinical observations, into a network, KGs provide a holistic representation of biomedical knowledge [23, 42, 50, 52]. Graph neural networks (GNNs) trained on such KGs can learn relational representations that enable data-driven prediction of drug indications [15, 57].

A notable recent advancement in this realm is TxGNN, a graph foundation model for clinician-centered drug repurposing [25]. TxGNN was trained on PrimeKG, a large-scale precision medicine KG that integrates over 20 data sources on diseases, drugs, genes, pathways, and more, and it demonstrated the ability to predict drug indications for over 17,000 diseases. TxGNN achieved substantial performance gains over previous methods[25]. This approach highlights how graph foundation models can transfer knowledge across diverse diseases to identify therapeutic candidates. PrimeKG’s rich collection of drug–disease and mechanistic relationships was central to TxGNN’s success, underscoring the importance of a high-quality, comprehensive KG in enabling powerful predictions.

Despite these advances, current GNN-based drug repurposing models have important limitations. In particular, few existing approaches integrate cell-type-specific molecular context into their KGs. Most biomedical KGs are not cell-type aware and do not capture how disease mechanisms or drug effects vary across different cell types. This limitation is especially critical for complex immune-related disorders such as autoimmune diseases, where the interplay between genetic risk loci and immune cell function is a key determinant of disease onset and progression [16, 35]. Autoimmune diseases represent a major area of unmet medical need and a compelling domain for repurposing, given their chronic nature and the overlapping immunological pathways involved in different conditions. Notably, studies have shown that many genetic influences on autoimmune disorders are highly cell-type specific: a variant may alter gene expression or function in one immune cell subtype (e.g., T-cells) while having little effect on others [16]. However, current KGs such as PrimeKG rarely include single-cell–resolution data, limiting their ability to capture links among genes, cell types, and diseases. As a result, gene–disease and drug–target relationships are averaged across tissues, preventing models from identifying which cell types drive disease or are therapeutically targeted.

Recent biomedical studies highlight the value of single-cell genomics in uncovering cell-type–specific disease mechanisms. For example, the OneK1K cohort shows that genetic variants can affect gene expression in one immune cell type but not in others [49, 54]. Such findings reinforce that the molecular context of a disease is tied to specific cell populations. We hypothesize that integrating these cell-type-specific expression into a KG could enrich a model’s understanding of disease biology and thereby improve its ability to predict effective drug–disease matches in the cellular context. GNNs have been applied to single-cell data for tasks such as modeling cell–cell networks and predicting drug–target relationships [30]. In contrast, KGs used for drug–disease repurposing are typically constructed without single-cell information, limiting their ability to capture cell-type–specific disease mechanisms and drug effects. Another limitation concerns how GNN-based repurposing models are evaluated. Prior studies often rely on randomly held-out drug–disease pairs, without considering coverage across the disease spectrum. As a result, test sets tend to be dominated by well-studied diseases, while diseases with sparse knowledge are underrepresented or excluded. This imbalance risks inflating performance metrics, because a model might appear accurate on average by mostly learning well-studied disease cases, yet fail to generalize to rarer or less-characterized diseases. Therefore, a more rigorous evaluation should ensure broad coverage of the disease spectrum. In this paper, we address the above limitations by making the following contributions: (i) We construct scPrimeKG, a reusable cell-aware knowledge graph: we curate and update PrimeKG into PrimeKG-U and enrich it with cell-type-specific expression signatures from the OneK1K cohort, adding cell-type resolution absent from existing biomedical KGs. (ii) We develop a GNN model, CellAwareGNN, that is pre-trained on scPrimeKG and then fine-tuned to predict drug–disease indications, with a focus on repurposing for autoimmune diseases. (iii) We introduce an evaluation benchmark with a per-disease stratified split that enables unbiased multi-KG comparison. Together, these contributions enable biologically informed drug repurposing by incorporating cell-type-specific expression context into an expanded knowledge graph, with CellAwareGNN trained under a pre-training and fine-tuning paradigm on scPrimeKG.

## 2 Background and Related Work

### 2.1 Biomedical knowledge graph

Computational approaches for drug repurposing often use KGs to represent biomedical entities as nodes and their interactions as edges [36]. These KGs integrate diverse data types from multiple sources into a single, unified graph structure. Homogeneous KGs contain only one type of node and edge. For example, protein– protein interaction networks represent molecular interactions and cellular signaling, which are critical for identifying potential drug targets [41]. In contrast, heterogeneous KGs incorporate multiple node and relation types to provide a more holistic view of biological systems. Recent studies include Bioteque, which integrates over 150 data sources to represent more than 450,000 biological entities across 12 node types (e.g., genes, diseases, and drugs) linked by 67 edge types [17]. Similarly, the OREGANO knowledge graph uses 12 node types and 19 edge types, with an emphasis on natural compounds such as herbal and plant remedies [10].

Among existing biomedical KGs, PrimeKG focuses on disease– drug associations. It includes 17,080 diseases with over 4 million relationships [12]. PrimeKG explicitly includes `indication, contra indication`, and `off-label` use relationships as drug–disease edges.

However, current KGs generally lack cell-type-specific details. Most genetic risks for complex diseases, especially autoimmune disorders, are mediated through specific cell types. Without cell-type-level resolution, regulatory mechanisms that occur only in particular cell types can be missed [16, 35]. To address this limitation, it is critical to integrate single-cell data (e.g., single-cell expression profiles) into KGs to better capture the cellular context needed for computational drug repurposing and precision medicine.

### 2.2 GNN for drug repurposing

Graph structures in the biomedical domain are complex and largescale, which requires effective representation learning to map biological entities into an embedding space. GNNs are a family of deep learning models designed to capture complex topological relationships between nodes (e.g., genes, pathways, and drugs). They can learn expressive representations of nodes, edges, and entire graphs. Therefore, drug repurposing is often defined as a link prediction problem, where GNNs learn embeddings that represent drugs, diseases, and their interactions by aggregating information from neighboring nodes.

Before GNN became prevalent, network-based methods were widely used to identify drug repurposing candidates by leveraging similarity networks and path-based inference on biomedical KGs [47]. For example, Chiang and Butte used a network-based method to generate 156,279 novel drug uses from 5,549 disease pairs that share at least one FDA-approved drug [13]. Many of these hypotheses are supported by clinical trials. Himmelstein et al. integrate heterogeneous biomedical knowledge from 29 data sources and use interpretable network features (Metapaths) to prioritize compound– disease pairs [24].

Recent literature has introduced several state-of-the-art (SOTA) GNN models for drug repurposing [26, 34, 56]. Sosa et al. proposed a framework that generates drug repurposing hypotheses for rare diseases using a knowledge graph constructed from literature-derived relationships [45]. TxGNN established SOTA performance for zero-shot drug repurposing, demonstrating promising predictions for diseases with limited treatment options [25]. DREAM-GNN learns representations from two complementary graphs (i.e., a known drug–disease interaction graph and a drug/disease similarity graph derived from chemical characteristics and protein sequences) [58]. Despite these advances, existing GNN models are limited in their ability to distinguish between a drug targeting a gene in different cellular states, such as resting T cells versus activated T cells. This lack of granularity can lead to suboptimal or incorrect predictions for diseases whose mechanisms depend on specific cellular contexts. Moreover, standard evaluation strategies for GNNs often sample edges uniformly at random, which is biased toward diseases and drugs with higher connectivity (i.e., those with more known treatments). This may allow models to achieve high performance by learning “shortcuts” based on the popularity of certain hub nodes, rather than learning biological principles that drive disease mechanisms.

We introduce CellAwareGNN to address these limitations by augmenting PrimeKG with single-cell expression context from the OneK1K cohort and using a disease-representation-aware evaluation framework.

## 3 Method

### 3.1 Problem definition

We model the biomedical domain as a heterogeneous knowledge graph *G* = (*V*,*E*,*R*), where *V* is the set of entities (e.g., drugs, diseases, genes, and cell types), *R* is the set of relation types (e.g., `indication`, `contraindication`, `causes`, and `associated_with`), and *E* ⊆ *V*×*R*×*V* is the set of typed edges. Each edge (*u, r, v*) ∈ *E* denotes that entity *u* is connected to entity *v* by relation *r* ∈ *R*.

Our goal is drug–disease link prediction for *indication*, which estimates the probability of an edge (*d*, `indication`, *s*) between a drug *d* and a disease *s*.

### 3.2 Single-cell knowledge graph construction

#### 3.2.1 Foundational Framework and Preprocessing (PrimeKG)

We utilize the PrimeKG framework [12] as the structural basis for our knowledge graph. PrimeKG is a multimodal network that integrates 20 primary high-quality data resources, bridging the gap between molecular heterogeneity and clinical phenotypes. The graph encompasses 10 distinct node types—including diseases, drugs, genes/proteins, phenotypes, pathways, and anatomical regions— across biological scales. Crucially for drug repurposing, it includes dense, typed therapeutic relationships, such as drug indications, contraindications, and off-label uses, sourced from DrugBank[27] and DrugCentral[48]. The construction and preprocessing of PrimeKG involve three rigorous standardization steps:

##### (1) Ontology Harmonization

We map all entities to standardized ontologies to ensure unique identification. For instance, drugs are mapped to DrugBank identifiers, diseases to the MONDO Disease Ontology [5], genes to NCBI Entrez IDs, and phenotypes to the Human Phenotype Ontology (HPO) [21].

##### (2) Overlap Resolution

To prevent node duplication, we resolve overlaps between ontologies. Specifically, phenotype nodes in HPO that share identifiers with disease nodes in MONDO are consolidated into single disease nodes, ensuring a clean separation between clinical diseases and phenotypic traits.

##### (3) Graph Cleaning

We merge the harmonized resources and extract the largest connected component to eliminate isolated sub-graphs, ensuring that all nodes are reachable during graph propagation.

#### 3.2.2 Temporal Update of Biomedical Knowledge (PrimeKG-U)

To ensure our model learns from the most current biomedical evidence, we construct PrimeKG-U. Instead of using the original 2021 data snapshots, we re-acquire and process all 20 primary data resources using their latest versions available as of November 2025. This update process involves re-executing the standardization pipeline on the newest releases of key databases, including DrugBank (v6.0+), DisGeNET [37] (v8.0), and the updated MONDO hierarchy. This results in a graph structure that retains the rigorous schema of PrimeKG but is populated with contemporary drug approvals and recently discovered gene-disease associations.

#### 3.2.3 Integration of Single-Cell Genomics (scPrimeKG)

We further extend PrimeKG-U to construct scPrimeKG, a single-cell enhanced knowledge graph that captures cell-type specific disease mechanisms. We integrate the OneK1K cohort dataset [54], which provides high-resolution single-cell RNA sequencing data from 1.27 million peripheral blood mononuclear cells collected from 982 healthy donors. The OneK1K study profiled single-cell gene expression across a broad range of immune cell populations, revealing extensive cell-type-specific transcriptional signatures. This integration introduces 22 immune cell type nodes (e.g., CD4+ T cells, NK cells, Monocytes) and gene-cell type edges to the graph structure.

We construct gene–cell type edges from cell-type-specific differential expression of the OneK1K single-cell transcriptomes. We build on the quality-controlled OneK1K data, which underwent filtering of low-quality cells, removal of lowly expressed genes, normalization, and correction for technical variation as described by Yazar et al. [54]. Single-cell read counts are first collapsed into a pseudo-bulk count matrix by grouping cells by donor and annotated cell type, and the resulting expression values are normalized, log-transformed, and scaled. For each gene, we then evaluate differential expression within a target cell type against all other cell types using the Wilcoxon rank-sum test, and link a gene to a cell type node only when it shows significant activation in that cell type at a Bonferroni-corrected *p*-value *<* 0.05. These stringent statistical thresholds ensure that only high-confidence, cell-type-specific associations are encoded as edges. By layering this single-cell resolution on top of the updated clinical knowledge base, scPrimeKG allows the model to differentiate between global and cell-type-specific gene expression, a critical factor for precision medicine in immune-mediated diseases.

### 3.3 Model architecture

Figure 1 provides an overview of the CellAwareGNN framework, including KG construction, two-phase training, and downstream drug–disease indication inference.

**Figure 1:**
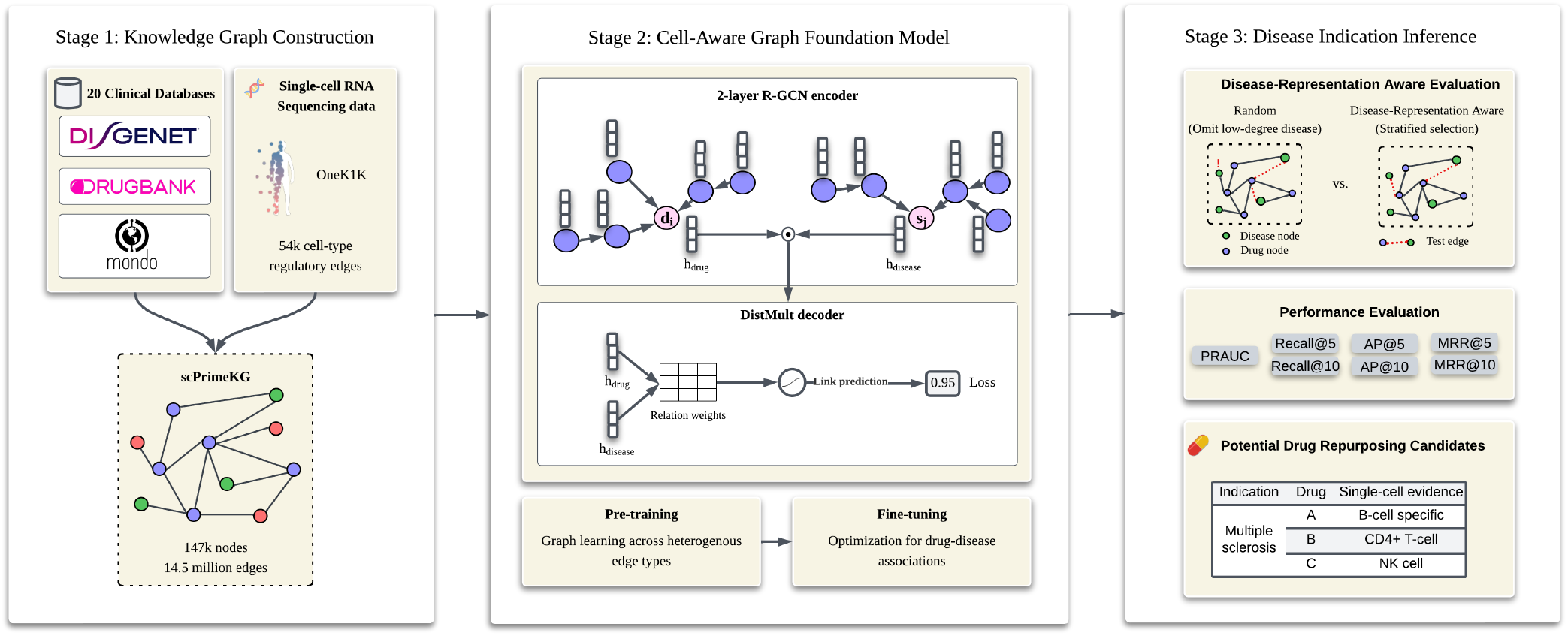
Overview of the CellAwareGNN Framework. The framework consists of three stages: (1) Knowledge Graph Construction, where standard clinical databases (e.g., DisGeNET, DrugBank, Mondo) are integrated with single-cell expression data (OneK1K) to construct scPrimeKG (14.5M edges); (2) Graph Foundation Model Training, utilizing a 2-layer R-GCN encoder and DistMult decoder with a two-phase strategy (general biological pre-training followed by drug–disease associations fine-tuning); and (3) Disease Indication Inference, where we implement a novel Disease-Representation Aware Evaluation strategy to rigorously assess generalization, with standard ranking metrics and the identification of drug repurposing candidates.

#### 3.3.1 CellAwareGNN overview

We use a two-layer heterogeneous relational graph convolutional network (R-GCN) as the encoder *f*_*θ*_ as in TxGNN. Each node embedding is initialized with Xavier initialization. For each node *v*, the encoder performs relation-aware message passing:

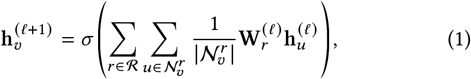

Where 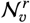denotes the neighbors of *v* under relation *r*, and 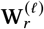is the relation-specific weight matrix at layer *ℓ*. The hidden size is 100.

The decoder is a relation-type aware DistMult [53]. For a query triplet (*u, r, v)*, we predict its likelihood using the DistMult scoring function:

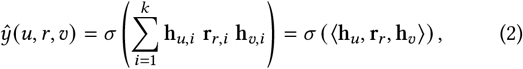

where *k* is the embedding dimension, h_*u*_, h_*v*_ ∈ ℝ^*k*^ are node embeddings, r_*r*∈_ ℝ^*k*^ is a learnable embedding for relation type *r*, and σ (·) is the sigmoid function.

We treat observed typed edges (*u, r, v*) ∈ E as positive training examples. For each positive triple (*u, r, v)*, we generate a negative triple (*u*^−^, *r, v*) by corrupting the source node while keeping the relation *r* and destination node *v* fixed. Specifically, we sample *u*^−^∈ V uniformly from entities that can appear as valid sources for relation *r* (i.e., entities with at least one outgoing edge of type *r*), and enforce *u*^−^ ≠ *u*.

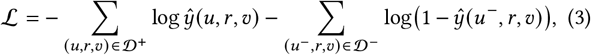

We optimize the model with a binary cross-entropy (BCE) loss over positive and negative triples:

where *D* ^+^ is the set of observed training triples and*D* ^−^ is the set of negatively sampled triples.

#### 3.2.2 Training pipeline

Training consists of two phases. In the pretrain phase, we update the encoder weights using all edge types in the KG. We use minibatch sampling and compute BCE loss across all relation types (batch size 1024, 2 epochs, learning rate 10^−3^). The goal is to learn a general graph representation from the full graph structure.

In the fine-tuning phase, we train on only drug–disease edges (i.e., indication, contraindication, and off-label use). We include contraindication and off-label use edges during training because they provide complementary real-world clinical signals: contraindications indicate safety and risk (i.e., relationships where a drug should be avoided for a disease), and off-label use edges reflect clinical practice patterns beyond approved labeling. Incorporating these additional drug–disease relations provides additional learning signal about drug–disease associations.

However, our evaluation focuses exclusively on indication predictions. Off-label use can introduce noise and variability because drugs may be prescribed outside their approved indications for heterogeneous reasons. Evaluating off-label use and contraindication prediction would require different ground-truth standards and is beyond the scope of this work.

We compute BCE loss on these drug–disease relation types only, focusing on learning drug–disease predictions (500 epochs, learning rate 5 ×10^−4^, following the same hyperparameters as TxGNN) [25]. The DistMult decoder weights are re-initialized before fine-tuning so that the model focuses on learning drug–disease specific relation representations.

#### 3.3.3 Clinical foundation model

We use the same model architecture across all models, we refer to the trained models as:

- **TxGNN**: TxGNN architecture trained on PrimeKG.
- **TxGNN-U**: the same architecture trained on PrimeKG-U.
- **CellAwareGNN**: the same architecture trained on scPrimeKG (single-cell genomics augmented KG).

## 4 Experiments

### 4.1 Experimental setup

#### Data splitting strategy

We implement disease-representation aware splitting to ensure representative evaluation across the diseases. For each disease with at least one drug–disease edge, we sample 30% of its edges for the test set, with a minimum of one edge per disease. This guarantees that every disease is represented in the test set (including rare diseases) and that each disease contributes proportionally to evaluation.

To enable consistent cross-KG evaluation, we use the same test split across TxGNN, TxGNN-U, and CellAwareGNN models evaluation. We repeat the splitting procedure across five random seeds.

To prevent label leakage, we remove inverse edges from training for all test-set drug–disease relationships. Knowledge graphs often store edges in both directions (e.g., Aspirin –indication–> Headache along with its reverse Headache –rev_indication–> Aspirin). If one of these reverse edges appears during training, the model can trivially predict test edges by memorizing the reverse relation rather than learning meaningful drug–disease associations. To avoid this, we ensure that for each indication edge in the test set, its reverse is excluded from training.

#### Autoimmune disease subset

From the test diseases, we isolate autoimmune diseases for focused evaluation. We defined the autoimmune subset using a curated disease list based on standard disease ontologies, following the classification established in the National Academies of Sciences, Engineering, and Medicine 2022 report [33].

#### Evaluation metrics

For each disease in the test set, we construct a candidate set consisting of all positive drug–disease pairs (known therapeutic relationships) and randomly sampled negative pairs at a 1:1 ratio: for a disease with *k* positive drugs, we randomly sample *k* negative drugs. This follows the evaluation methodology used in TxGNN and yields more meaningful discrimination metrics by focusing on harder negatives.

On the balanced candidate set, we compute AUPRC. Additionally, we report Recall@*k*, Average Precision@*k* (AP@*k*), and Mean Reciprocal Rank@*k* (MRR@*k*) at *k* ∈ {5, 10, 30}. For MRR@*k*, we compute the reciprocal rank 1 rank if the first ground-truth drug appears within the top *k* predictions and 0 otherwise. We report results for *k* ∈ {5, 10} in the main text and include *k* = 30 in the Appendix.

We report overall metrics averaged across all test diseases and the metrics computed over the autoimmune disease subset. All results are reported as mean and 95% confidence intervals (CI) across five random seeds.

### 4.2 Knowledge Graph Statistics

Table 1 summarizes the size and composition of the knowledge graphs used in this study. PrimeKG-U is an updated biomedical baseline that expands PrimeKG coverage by incorporating newer releases of key biomedical databases, increasing the number of edges from 8.10M to 14.47M (+78%) and the number of nodes from 129,312 to 147,859. Our key novelty is scPrimeKG, a single-cell–enhanced knowledge graph that extends PrimeKG-U with cell-type–resolved expression evidence while keeping the overall graph scale comparable (147,881 nodes and 14.52M edges). Specifically, scPrimeKG introduces 22 cell-type nodes and 54,670 gene-cell type edges that capture cell-type-specific gene expression, providing fine-grained, cell-aware genomic context that is absent from existing biomedical KGs. The single-cell augmentation adds only 0.38% to the edge count and incurs negligible training overhead (Appendix D).

**Table 1:** Comparison of knowledge graph statistics. PrimeKG-U is the updated biomedical baseline (+78% edges over original PrimeKG). scPrimeKG (ours) further integrates cell-type nodes and gene–cell type edges capturing cell-type-specific expression.

| Statistic | PrimeKG (Original) | PrimeKG-U (Updated) | scPrimeKG (Ours) |
| --- | --- | --- | --- |
| *Overall* |  |  |  |
| Total Nodes | 129,312 | 147,859 | 147,881 |
| Total Edges | 8,100,498 | 14,472,538 | 14,527,208 |
| *Node types* |  |  |  |
| Gene/Protein | 27,671 | 35,787 | 35,787 |
| Biological Process | 28,642 | 29,959 | 29,959 |
| Effect/Phenotype | 15,311 | 19,362 | 19,362 |
| Diseases | 17,080 | 18,071 | 18,071 |
| Anatomy | 14,035 | 14,638 | 14,638 |
| Molecular Function | 11,169 | 12,509 | 12,509 |
| Drug | 7,957 | 9,503 | 9,503 |
| Cellular Component | 4,176 | 4,309 | 4,309 |
| Pathway | 2,516 | 2,834 | 2,834 |
| Exposure | 818 | 887 | 887 |
| Cell types | – | – | 22 |
| *Edge types* |  |  |  |
| Anatomy-Protein(Present) | 3,036,406 | 7,958,182 | 7,958,182 |
| Drug-Drug | 2,672,628 | 3,061,974 | 3,061,974 |
| Anatomy-Protein(Absent) | 39,774 | 744,214 | 744,214 |
| Protein-Protein | 642,150 | 648,124 | 648,124 |
| Disease-Phenotype(Positive) | 300,634 | 398,504 | 398,504 |
| Biological process-Protein | 289,610 | 360,656 | 360,656 |
| Cellular component-Protein | 166,804 | 194,740 | 194,740 |
| Molecular function-Protein | 139,060 | 180,782 | 180,782 |
| Disease-Protein | 160,822 | 172,030 | 172,030 |
| Drug-Effect | 129,568 | 132,716 | 132,716 |
| Biological process-Biological process | 105,772 | 117,546 | 117,546 |
| Pathway-Protein | 85,292 | 94,262 | 94,262 |
| Disease-Disease | 64,388 | 82,680 | 82,680 |
| Contraindication | 61,350 | 71,946 | 71,946 |
| Drug-Protein | 51,306 | 63,900 | 63,900 |
| Phenotype-Phenotype | 37,472 | 50,332 | 50,332 |
| Molecular function-Molecular function | 27,148 | 32,848 | 32,848 |
| Anatomy-Anatomy | 28,064 | 31,364 | 31,364 |
| Indication | 18,776 | 21,742 | 21,742 |
| Cellular component-Cellular component | 9,690 | 11,396 | 11,396 |
| Phenotype-Protein | 6,660 | 7,096 | 7,096 |
| Exposure-Disease | 4,608 | 6,282 | 6,282 |
| Pathway-Pathway | 5,070 | 5,852 | 5,852 |
| Off-Label-Use | 5,136 | 5,750 | 5,750 |
| Exposure-Protein | 2,424 | 5,250 | 5,250 |
| Exposure-Exposure | 4,140 | 4,972 | 4,972 |
| Exposure-Biological process | 3,250 | 4,488 | 4,488 |
| Disease-Phenotype(Negative) | 2,386 | 2,772 | 2,772 |
| Exposure-Molecular function | 90 | 112 | 112 |
| Exposure-Cellular component | 20 | 26 | 26 |
| Protein-Cell type | – | – | 54,670 |

### 4.3 Overall drug retrieval performance

The test set comprises 1,169 unique diseases, with an average of 2.8 ± 3.2 indication edges per disease. This reflects that most diseases have only 1 to 3 known drug indications in the test set, while a smaller number of well-characterized diseases have larger numbers of treatment associations.

As shown in Table 2, we observe consistent improvements in model performance. First, updating the KG (TxGNN → TxGNN-U) improves overall indication retrieval, increasing AUPRC from 0.799 to 0.816 and improving top-*k* retrieval at both *k* = 5 and *k* = 10 (e.g., Recall@5: 0.902 to 0.907 and Recall@10: 0.969 to 0.973; AP@5: 0.823 to 0.836 and AP@10: 0.807 to 0.822; MRR@5: 0.841 to 0.854 and MRR@10: 0.841 to 0.854).

**Table 2:** Indication prediction evaluation (mean [95% CI]).

| Model | AUPRC | Recall@5 | Recall@10 | AP@5 | AP@10 | MRR@5 | MRR@10 |
| --- | --- | --- | --- | --- | --- | --- | --- |
| TxGNN | 0.7990 [0.7904, 0.8075] | 0.9020 [0.8995, 0.9045] | 0.9691 [0.9667, 0.9714] | 0.8228 [0.8133, 0.8323] | 0.8071 [0.7982, 0.8159] | 0.8413 [0.8330, 0.8497] | 0.8413 [0.8330, 0.8497] |
| TxGNN-U | 0.8156 [0.8095, 0.8217] | 0.9070 [0.9034, 0.9106] | 0.9730 [0.9714, 0.9746] | 0.8364 [0.8285, 0.8443] | 0.8224 [0.8162, 0.8287] | 0.8538 [0.8456, 0.8621] | 0.8538 [0.8456, 0.8621] |
| CellAwareGNN | **0.8264** [0.8181, 0.8347] | **0.9101** [0.9074, 0.9127] | **0.9751** [0.9733, 0.9769] | **0.8462** [0.8378, 0.8547] | **0.8328** [0.8250, 0.8406] | **0.8636** [0.8564, 0.8708] | **0.8636** [0.8564, 0.8708] |

We then evaluated whether incorporating single-cell sequencing data further improves performance. On the overall test set, CellAwareGNN achieves the best performance across all reported metrics. Specifically, CellAwareGNN achieves an AUPRC of 0.826, a Recall@5 of 0.910 and Recall@10 of 0.975, an AP@5 of 0.846 and AP@10 of 0.833, and an MRR@5 and MRR@10 of 0.864. These results indicate that integrating cell-type edges improves overall retrieval quality.

### 4.4 Domain-specific drug retrieval

We further evaluate performance on an autoimmune disease subset of the test set, which contains 40 autoimmune diseases with an average of 3.2 ± 3.5 indication edges per disease. Compared to the overall test set, autoimmune diseases have relatively more treatments approved.

Table 3 summarizes the results. Updating the underlying databases (TxGNN → TxGNN-U) improves AUPRC from 0.815 to 0.847 and increases top-*k* retrieval quality at both *k* = 5 and *k* = 10. Specifically, TxGNN-U improves Recall@5 from 0.885 to 0.896 and Recall@10 from 0.956 to 0.965, improves AP@5 from 0.851 to 0.879 and AP@10 from 0.828 to 0.864, and improves MRR@5 and MRR@10 from 0.863 to 0.887.

**Table 3:** Autoimmune indication prediction evaluation (mean [95% CI]).

| Model | AUPRC | Recall@5 | Recall@10 | AP@5 | AP@10 | MRR@5 | MRR@10 |
| --- | --- | --- | --- | --- | --- | --- | --- |
| TxGNN | 0.8147 [0.7914, 0.8379] | 0.8854 [0.8738, 0.8969] | 0.9562 [0.9482, 0.9641] | 0.8505 [0.8290, 0.8720] | 0.8283 [0.8070, 0.8495] | 0.8631 [0.8329, 0.8933] | 0.8631 [0.8329, 0.8933] |
| TxGNN-U | 0.8466 [0.8240, 0.8692] | 0.8955 [0.8857, 0.9054] | 0.9649 [0.9567, 0.9730] | 0.8790 [0.8585, 0.8995] | 0.8643 [0.8439, 0.8847] | 0.8871 [0.8664, 0.9079] | 0.8871 [0.8664, 0.9079] |
| CellAwareGNN | **0.8636** [0.8476, 0.8797] | **0.8961** [0.8824, 0.9098] | **0.9707** [0.9617, 0.9798] | **0.8962** [0.8852, 0.9073] | **0.8776** [0.8656, 0.8896] | **0.9068** [0.8911, 0.9226] | **0.9068** [0.8911, 0.9226] |

Incorporating single-cell data with CellAwareGNN further improves retrieval across all reported metrics. CellAwareGNN achieves the best AUPRC (0.864), as well as the best ranking performance, with a Recall@5 of 0.896 and Recall@10 of 0.971, an AP@5 of 0.896 and AP@10 of 0.878, and an MRR@5 and MRR@10 of 0.907. Overall, CellAwareGNN consistently outperforms the TxGNN-U baseline on autoimmune indications.

### 4.5 Qualitative analysis: top ranked drugs validation and discovery

In this section, we provide qualitative evidence that CellAwareGNN’s improvements are not limited to aggregate metrics. We organize the analysis into (i) *KG-novel* repurposing candidates (edges absent from the scPrimeKG training graph) supported by single-cell mechanistic rationale and emerging clinical evidence, and (ii) validation against standard-of-care therapies.

#### 4.5.1 KG-novel candidates

##### Pemphigus Novel Repurposing Candidates

Pemphigus is a rare autoimmune blistering disease affecting the skin and mucous membranes. CellAwareGNN ranks ocrelizumab (#22) and methotrexate (#19) as novel candidates, which are supported by the single-cell evidence in scPrimeKG. Ocrelizumab targets CD20 (MS4A1), which OneK1K data shows is highly expressed in B naive, B intermediate, and B memory cells [54]. These B cell populations give rise to autoreactive plasma cells that produce pathogenic anti-desmoglein autoantibodies in Pemphigus [38, 39]. Since rituximab (another antiCD20 therapy) is FDA-approved for Pemphigus [20], ocrelizumab represents a clinically plausible alternative. A case study reported successful treatment of a patient with 5-year history of refractory Pemphigus using ocrelizumab (two 300 mg infusions given 2 weeks apart per cycle) following a rituximab allergy [9]. Methotrexate targets DHFR and ATIC, genes expressed across proliferating T and B cells in OneK1K. It reduces inflammation and modulates immune function, leading to disease suppression. Current clinical data suggests that methotrexate is a useful steroid-sparing agent, particularly in cases of steroid-dependent Pemphigus with ongoing trials and studies evaluating the efficacy of methotrexate for Pemphigus [8, 46].

##### Rheumatoid Arthritis (RA) Novel Repurposing Candidates

For RA, CellAwareGNN identifies emapalumab (#26) and rosiglitazone (#29) as novel candidates with strong single-cell rationale. Emapalumab targets IFN-γ, which is a key cytokine involved in inflammatory pathways in RA [14, 43]. The EMERALD study (NCT05001737), an open-label trial, evaluates emapalumab in pediatric and adult patients with macrophage activation syndrome (MAS) associated with two systemic inflammatory diseases, Still’s disease or Systemic Lupus Erythematosus, supporting its potential in RA [11]. Rosiglitazone activates PPAR-γ, which has anti-inflammatory properties including downregulation of pro-inflammatory cytokines such TNF-*α*, IL-6, and IL-1 [31]. This suggests that it could theoretically be beneficial in managing inflammation in diseases like RA. A completed clinical trial (NCT00379600) evaluated rosiglitazone’s anti-inflammatory and metabolic effects in RA subjects, though results are not published yet [22].

#### 4.5.2 Validation against standard-of-care therapies

##### Pemphigus validation

To validate that single-cell expression data in scPrimeKG improves drug ranking performance, we compare CellAwareGNN predictions against TxGNN-U baseline for known Pemphigus treatments. CellAwareGNN ranks prednisolone at Rank #1, which is consistent with expert recommendations that systemic corticosteroids (e.g., prednisone/prednisolone) are a first-line treatment for Pemphigus [1, 32]. Azathioprine is a first-line immunosuppressant medication for Pemphigus [38]. CellAwareGNN ranks it at #12, which TxGNN-U ranks at #22. Azathioprine works by inhibiting intracellular purine synthesis in rapidly dividing cells, which reduces the proliferation of circulating T and B lymphocytes [2]. This improvement demonstrates CellAwareGNN’s ability to correctly identify azathioprine’s mechanism of directly targeting the autoreactive cell populations in Pemphigus. CellAwareGNN also improved the rank of methylprednisolone, a corticosteroid that treats severe Pemphigus, from rank #9 to #6 and moved down the rank of hydrocortisone from #3 to #7. These results align with in vitro studies showing that methylprednisolone is a more potent lymphocyte suppressor than hydrocortisone [29]. By leveraging OneK1K data, our model successfully captures the cell-type-specific gene expression that drives this difference in potency.

##### Multiple sclerosis (MS) validation

For MS, CellAwareGNN correctly prioritizes methylprednisolone (from #14 predicted by TxGNN-U to #8), consistent with clinical guidelines that using high-dose methylprednisolone as standard of care for treating acute MS relapses [4, 7]. MS is driven by specific immune subpopulations, T-cells and B-cells subsets, that infiltrate the nervous system [3]. By integrating single-cell expression data from OneK1K, CellAwareGNN explicitly models the high expression of glucocorticoid receptors within these pathogenic T and B cell subsets. This allows the model to computationally link the methylprednisolone’s mechanism to the MS’s specific cellular drivers. Notably, CellAwareGNN also improves the ranking of ocrelizumab from #30 to #20. It is a FDA-approved anti-CD20 therapy. This improvement aligns with single-cell studies showing that ocrelizumab binds to CD20+ B cells, a cell type that contributes to nerve damage in MS patients, and therefore reduces the immune system’s attack on the nervous system [6, 51]. These results demonstrate that single-cell expression integration enables CellAwareGNN to capture both glucocorticoid potency differences and the emerging importance of B cell targeting in MS treatment.

##### Rheumatoid arthritis (RA) validation

TxGNN-U’s predictions are dominated by corticosteroids (e.g., prednisone and triamcinolone). In contrast, CellAwareGNN diversifies the candidate list by retrieving disease-modifying antirheumatic drugs (DMARDs). Azathioprine increases from Rank #12 by TxGNN-U to #7, which targets CD4+ T cells identified in RA synovium [19, 55]. These results suggest that CellAwareGNN better captures cell-type-specific drug target expression patterns that are relevant to the heterogeneous immune cell populations driving RA inflammation.

### 4.6 Quantifying the Contribution of Cell-Type Edges

To quantify the relative importance of cell-type edges compared to conventional biomedical KG edges, we construct variants of CellAwareGNN by removing individual edge types (Drug–Protein, Drug–Drug, Drug–Effect, Disease–Protein, or Disease–Phenotype) and compare their performance against TxGNN-U, which contains all other biomedical KG edges but lacks cell-type edges.

- **CellAwareGNN-DP (No Drug–Protein):** We remove all drug_protein edges (drug–gene associations) while retaining all other edge types. This tests whether drug–gene relationships contribute to prediction performance.
- **CellAwareGNN-DD (No Drug–Drug):** We remove all drug– drug edges (drug–drug interactions) while retaining all other edge types. This tests whether drug–drug connectivity contributes to prediction performance.
- **CellAwareGNN-DPh (No Drug–Effect):** We remove all drug–effect edges while retaining all other edge types. This tests whether phenotypic relationships contribute to prediction performance.
- **CellAwareGNN-SP (No Disease–Protein):** We remove all disease_protein edges (disease–gene associations) while retaining all other edge types. This tests whether disease– gene relationships contribute to prediction performance.
- **CellAwareGNN-SPh (No Disease–Phenotype):** We remove all disease–phenotype-positive edges (disease–phenotype positive associations) while retaining all other edge types. This tests whether phenotype annotations for diseases contribute to prediction performance.

All CellAwareGNN variants are trained with the same hyperparameters and evaluated on the same disease-stratified test set.

Figure 2 summarizes the results for the overall and autoimmune disease subsets. We make two observations. First, AUPRC and ranking metrics at *k* = 10 remain stable across variants: removing any single biomedical edge type leads to only modest changes in performance. Across variants, AUPRC remains consistently high (0.818– 0.824), Recall@10 is similar (0.966–0.970), AP@10 varies slightly (0.864–0.876), and MRR@10 is stable (0.856–0.867). This robustness indicates that when one edge type is removed, CellAwareGNN can leverage alternative relational paths for information flow and maintain accurate drug ranking. For example, without drug–protein edges, drug–disease signals can still be inferred through paths such as drug–effect/phenotype and phenotype–disease edges.

**Figure 2:**
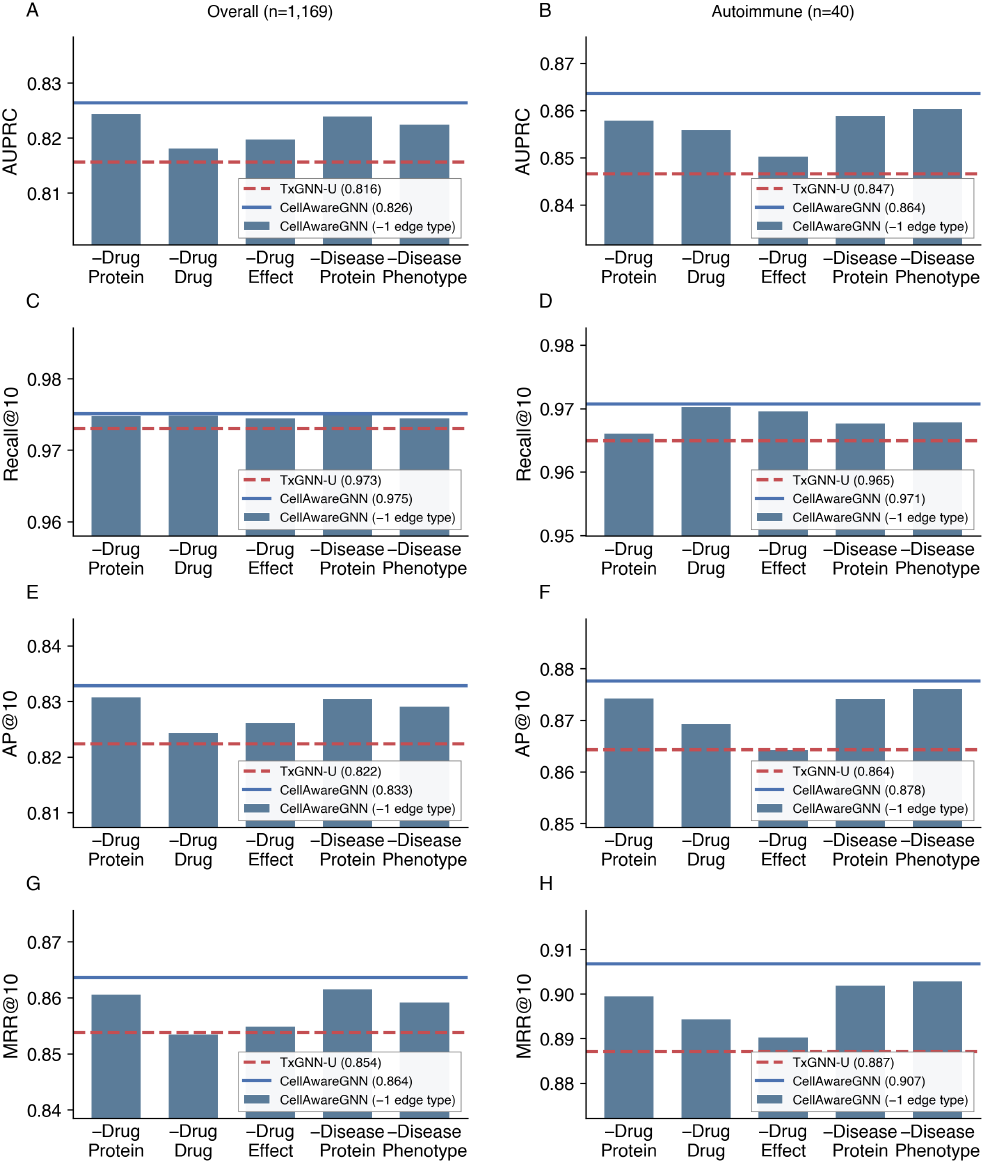
Contribution of cell-type edges compared to conventional KG edges. CellAwareGNN variants with one edge type removed are compared against TxGNN-U (which has all other edges but no cell-type edges).

Second, we observe that CellAwareGNN variants consistently outperform TxGNN-U on drug indication prediction in both overall disease test set and the autoimmune disease subset. For example, CellAwareGNN-DD (i.e., with drug–drug edges removed from scPrimeKG) achieves an AUPRC (95% CI) of 0.818 [0.807, 0.828] on the overall test set and 0.8559 [0.829, 0.882] on the autoimmune subset; the same trend is observed for other ranking metrics (Recall@10 = 0.974 [0.973, 0.976], AP@10 = 0.824 [0.813, 0.834], and MRR@10 = 0.853 [0.840, 0.866]).

To separate biological signal from graph-density effects, we additionally compared scPrimeKG against a control graph in which cell-type nodes are randomly connected to other nodes while preserving the number and degree distribution of cell-type edges (Appendix Table 7). Random connectivity recovers only a small fraction of the improvement over TxGNN-U (35% of the AUPRC gain for autoimmune diseases), with the majority attributable to biologically specific gene–cell-type edges, indicating that biological content drives the improvement rather than added graph density.

Overall, these findings suggest that the augmented cell-type edges are not redundant information beyond conventional drug, disease, and gene relations. They support our hypothesis that incorporating single-cell expression evidence into the KG enhances drug indication prediction, particularly for autoimmune diseases.

## 5 Computation Environment and Reproducibility

All experiments were conducted on two NVIDIA RTX A6000 GPUs using Python 3.8.20. We fix random seeds and use predefined data splits for training, validation, and testing.

To support reproducibility, we released training and evaluation scripts, configuration files, and data-processing pipeline at https://github.com/OHPENLab/CellAwareGNN.

## 6 Limitations and Ethical Considerations

This study does not involve newly-collected patient-level data. We use publicly available databases and adhere to the corresponding data-use agreements. The biomedical databases we include (e.g., DrugBank and Mondo) may reflect bias toward well-studied diseases, drugs, and genes, and such biases can propagate into model predictions.

Our study has several limitations. First, our evaluation focuses on ranking metrics and literature. Additional validation on real-world data such as electronic health records is needed to assess clinical utility. Second, our model learns on population-level KG and does not incorporate patient-level factors (e.g., individual variability in genes, environment, and lifestyle) that may influence individual treatment response. Third, this study focuses on the investigation of the contribution of single-cell data to drug indication prediction. Future study should explore model architectures that are more tailored to single-cell data incorporation. Fourth, our single-cell evidence is limited to peripheral blood mononuclear cells from the OneK1K cohort, so tissue-resident cell types are not represented. As a result, the autoimmune benefit is the primary validated use case, and generalization to other disease areas awaits integration of tissue-specific single-cell data. Relatedly, OneK1K comprises healthy donors, so scPrimeKG encodes constitutive rather than disease-perturbed expression; extending it with disease-cohort single-cell atlases warrants further investigation.

## 7 Conclusion

We introduced scPrimeKG, a single-cell–enhanced biomedical knowledge graph, and CellAwareGNN, a graph foundation model that incorporates cell-type–resolved expression evidence for drug–disease indication prediction. Our approach improves predictive performance across overall and autoimmune-focused evaluations and prioritizes biologically plausible repurposing hypotheses, such as Ocrelizumab and Methotrexate for Pemphigus and Rosiglitazone for Rheumatoid Arthritis. These candidates represent computationally inferred signals that warrant further validation using real-world data and experimental or clinical studies.

Looking forward, this work establishes a foundation for extending graph-based drug repurposing with richer cellular context. Future directions include expanding scPrimeKG to represent tissue context and cell–cell interactions, and developing R-GCN–based architectures that explicitly model hierarchical relationships among genes, cell types, and diseases within the knowledge graph. Together, these extensions may further improve the accuracy, interpretability, and translational relevance of cell-aware graph foundation models for drug repurposing.

## Acknowledgments

This work was supported in part by the National Library of Medicine of the National Institutes of Health under Award Numbers R01LM014199 and K99LM014428, and by the National Cancer Institute of the National Institutes of Health under Award Numbers T32CA160056 and R01CA297582.

## A. Experiments

### A.1. Additional Indication Prediction Results (*k* = 30)

Table 4 and Table 5 report indication prediction performance at *k* = 30 for the overall test set and the autoimmune disease subset, respectively. We include these metrics in addition to the main-text results at *k* ∈ {5, 10}.

**Table 4:** Indication prediction evaluation at *k* = 30 (mean [95% CI]).

| Model | AUPRC | Recall@30 | AP@30 | MRR@30 |
| --- | --- | --- | --- | --- |
| TxGNN | 0.7990 [0.7904, 0.8075] | 0.9979 [0.9976, 0.9982] | 0.7994 [0.7908, 0.8081] | 0.8413 [0.8330, 0.8497] |
| TxGNN-U | 0.8156 [0.8095, 0.8217] | 0.9981 [0.9978, 0.9985] | 0.8162 [0.8103, 0.8221] | 0.8538 [0.8456, 0.8621] |
| CellAwareGNN | **0.8264** [0.8181, 0.8347] | **0.9982** [0.9978, 0.9987] | **0.8271** [0.8193, 0.8350] | **0.8636** [0.8564, 0.8708] |

**Table 5:** Autoimmune indication prediction evaluation at *k* = 30 (mean [95% CI]).

| Model | AUPRC | Recall@30 | AP@30 | MRR@30 |
| --- | --- | --- | --- | --- |
| TxGNN | 0.8147 [0.7914, 0.8379] | 0.9942 [0.9930, 0.9953] | 0.8160 [0.7932, 0.8388] | 0.8631 [0.8329, 0.8933] |
| TxGNN-U | 0.8466 [0.8240, 0.8692] | 0.9942 [0.9922, 0.9962] | 0.8543 [0.8309, 0.8777] | 0.8871 [0.8664, 0.9079] |
| CellAwareGNN | **0.8636** [0.8476, 0.8797] | **0.9952** [0.9935, 0.9969] | **0.8698** [0.8554, 0.8842] | **0.9068** [0.8911, 0.9226] |

## B Top-Ranked Drug Candidates

Table 6 report the top-30 ranked drug candidates for several autoimmune diseases produced by CellAwareGNN. Bold entries denote KG-novel predictions (i.e., drug–disease edges that are not present in the training graph).

**Table 6:** Top-30 predicted drug candidates by CellAwareGNN. Bold text indicates KG-Novel candidates (edges not present in the training graph).

| Rank | Pemphigus | Rheumatoid Arthritis | Multiple sclerosis |
| --- | --- | --- | --- |
| 1 | Prednisolone | Prednisone | **Hydroxyprogesterone caproate** |
| 2 | Cortisone acetate | Methylprednisolone | **Difluprednate** |
| 3 | Dexamethasone | Triamcinolone | Hydrocortisone acetate |
| 4 | Betamethasone | Betamethasone | **Streptozocin** |
| 5 | Prednisone | Dexamethasone | **Tamoxifen** |
| 6 | Methylprednisolone | Azathioprine | **Stanolone** |
| 7 | Hydrocortisone | Hydrocortisone | Betamethasone |
| 8 | Triamcinolone | Hydrocortisone | Methylprednisolone |
| 9 | Hydrocortisone acetate | Hydrocortisone acetate | Prednisolone |
| 10 | **Difluprednate** | Cortisone acetate | Cortisone acetate |
| 11 | **Desoximetasone** | Sulfasalazine | **Nandrolone** |
| 12 | Azathioprine | Peficitinib | **Calcipotriol** |
| 13 | **Desonide** | **Acitretin** | **Anastrozole** |
| 14 | **Fluocinolone acetonide** | Methotrexate | **Ifosfamide** |
| 15 | **Stanolone** | Tenidap | Prednisone |
| 16 | **Indomethacin** | **Ifosfamide** | Dexamethasone |
| 17 | **Amcinonide** | **Ocrelizumab** | **Acitretin** |
| 18 | **Hydroxychloroquine** | **Oxandrolone** | **Oxaliplatin** |
| 19 | **Methotrexate** | **Hydroxyprogesterone caproate** | **Diethylstilbestrol** |
| 20 | **Flumethasone** | **Levonorgestrel** | **Ocrelizumab** |
| 21 | **Calcipotriol** | **Difluprednate** | Hydrocortisone |
| 22 | **Ocrelizumab** | Diclofenac | Triamcinolone |
| 23 | **Streptozocin** | **Baclofen** | **Chlorotrianisene** |
| 24 | **Chloroquine** | **Misoprostol** | **DB14776** |
| 25 | **Fluocinonide** | **Mesalazine** | **Aminoglutethimide** |
| 26 | **Diclofenac** | **Emapalumab** | **Mitotane** |
| 27 | **DB02046** | **Mitotane** | **Estetrol** |
| 28 | **Furoyl-Leucine** | **Streptozocin** | **Clomifene** |
| 29 | **2-(Thiomethylene)-4-methylpentanoic acid** | **Rosiglitazone** | **Etonogestrel** |
| 30 | **Rimexolone** | Sarilumab | **Lebrikizumab** |

## C. Random connection: Biological Signal vs. Graph Density

To test whether the performance gains arise from biologically meaningful cell-type content rather than from increased graph density, we constructed a control graph in which cell-type nodes are randomly connected to existing nodes, preserving both the number of cell-type edges and their degree distribution. All models were evaluated over 5 random seeds; we report average AUPRC (Table 7).

**Table 7:** AUPRC comparison across CellAwareGNN, random-connection control, and TxGNN-U variants. Computational Efficiency.

| Model | All | Autoimmune |
| --- | --- | --- |
| CellAwareGNN (scPrimeKG) | 0.826 | 0.864 |
| CellAwareGNN (Random) | 0.822 | 0.853 |
| TxGNN-U (PrimeKG-U) | 0.816 | 0.847 |

Graph density alone yields a marginal benefit: the random-connection variant improves only slightly over TxGNN-U (+0.006 AUPRC over-

all and autoimmune). The full scPrimeKG outperforms the random-connection variant across all metrics. In the autoimmune subset, random connectivity accounts for only 35% of the total gain over TxGNN-U (0.006 of 0.017 AUPRC), with the remaining 65% attributable to biologically specific gene–cell-type edges.

All three models were benchmarked under identical conditions (2 pre-training epochs + 500 fine-tuning epochs; single A6000 GPU). As shown in Table 8, CellAwareGNN had comparable runtime and GPU memory usage to TxGNN-U.

**Table 8:** Computational efficiency under identical bench-marking conditions.

| Model | Time (min) | Memory (MiB) |
| --- | --- | --- |
| TxGNN | 100.3 | 1,394 |
| TxGNN-U | 235.7 | 2,293 |
| CellAwareGNN | 237.8 | 2,330 |

